# Characterization of Novel Peptides with Antimicrobial and Immunomodulatory Potential

**DOI:** 10.1101/2023.07.07.548068

**Authors:** Swetha Kakkerla, Sridhar Kavela, Mohammad Kadivella, Murali Krishna Thupurani, Sathvika Chintalpani

## Abstract

In this study, we designed six peptides based on the characteristics of natural antimicrobial peptides (AMPs), including small size, cationicity, amphipathicity, and *α*-helix structure. The peptides were evaluated for their antimicrobial activity, hemolytic potential, binding affinity to TLR4/MD-2 receptors, and immunomodulatory effects. Five out of the six designed peptides were classified as AMPs, while one peptide was predicted to be a non-antimicrobial peptide (NAMP) by the AxPEP server. The designed peptides exhibited varying degrees of antimicrobial activity against Escherichia coli, Staphylococcus aureus, Klebsiella pneumoniae, and Pseudomonas aeruginosa, with SK1217 and SK1260 demonstrating stronger antimicrobial potential. Hemolytic assays revealed minimal hemolytic activity for SK1203, SK1217, SK1260, and SK1281, indicating a low propensity to cause red blood cell (RBC) lysis. However, SK1286 and SK1283 exhibited significant hemolytic effects at higher concentrations, with SK1286 causing complete hemolysis at 100 μg/mL. Molecular docking analysis showed favourable binding affinities of SK1260, SK1217, and SK1286 with TLR4/MD-2 receptors. Furthermore, SK1260 displayed potent immunomodulatory activity, reducing the release of pro-inflammatory cytokines IL-6 and TNF-α in LPS-stimulated macrophage cells. Cell viability assays demonstrated minimal cytotoxicity of SK1260. In vivo studies on infected mice showed that SK1260 treatment increased leukocyte recruitment and cytokine levels (GM-CSF, INF-g, and MCP-1) in the peritoneal lavage fluid. Overall, these findings highlight the promising potential of these cationic AMPs as antimicrobial and immunomodulatory agents for further therapeutic exploration.

## 1. Introduction

Drug resistance to microorganisms, such as fungi, viruses, and parasites as well as bacteria like staphylococci, enterococci, and Escherichia coli, has grown to be a significant global health issue. Since most living things naturally produce antimicrobial peptides (AMPs) to fend against encroaching pathogens, they make excellent candidates for development as novel antibiotics [M. Zasloff et al. 2002, R. E. W. Hancock et al. 2002 and J. M. Thomson et al. 2005]. Several of them have also been discovered to have spermicidal and anticancer properties [R. Lai et al. 2002 and M.A Baker et al. 1993]. These AMPs are cationic, amphipathic, and only slightly larger than 10 kDa in MW, with varying lengths, sequences, and structures. The first Antimicrobial peptides were discovered in a giant silk moth Hyalophora cecropin in 1981 (H. Steiner et al. 1981). Peptides with antibacterial action have since been discovered in all members of the natural world, including bacteria, insects, amphibians, and mammals [W. F. Broekaert et al. 1997, S. Haeberli et al. 2000 and H.G Boman et al. 1991]. To exert their antimicrobial effect, they employ a typical “carpet” method known as accumulation on the bacterial membrane up to a threshold concentration, which leads to membrane permeabilization/disintegration [Y. Shai et al. 2001].

The AMPs are divided into five groups [K. V. R. Reddy et al. 2004]: (1) peptides that form alpha-helical structures, (2) peptides rich in cysteine residues, (3) peptides that form beta-sheet structures, (4) peptides rich in regular amino acids, such as histidine, arginine, and proline, and (5) peptides made up of uncommon and modified amino acids. The design and synthesis of potent AMPs can be guided by these general characteristics.

The AMPs that exist in nature are far from optimal, and a few of these can be toxic to eukaryotes. For instance, the hemolysis of eukaryotes is commonly accompanied by strong antibacterial action. The primary goal of the study is to improve their antibacterial effectiveness and decrease their toxic effects on eukaryotes. In order to overcome the limitations in large-scale manufacturing and application of natural peptides, such as the low specific activity and the limited amounts recovered from natural sources, rational design and subsequent chemical synthesis are crucial approaches. For designing active peptides, there are several databases and programs available. The website http://web.expasy.org/blast/ is a helpful resource for secondary structure prediction and sequence analysis. Strong functionality is offered by Accelrys Insight II for investigating the structure and target-based drug discovery. It is known that AMPs’ physical and chemical characteristics, such as secondary structure, total charge, and hydrophobicity, influence how well they interact with model membranes and mammalian cells [H.-S. Ahn et al. 2006, H.-T. Chou, et al. 2008, H. Jenssen et al. 2008 and Giuliani, G. Pirri et al. 2007]. As AMPs interact with the target microorganism, the majority of them fold into an amphipathic-helical shape. Designing AMPs that are more effective at killing bacteria and less hazardous to eukaryotes is now possible thanks to advancements in structure-activity relationship and mechanism studies. Both naturally occurring peptides and synthetic ones have been shown to have potent antimicrobial, anti-biofilm, and immunomodulatory properties (de la Fuente-Núñez et al, 2014, Hilchie, A. L et al 2013 & Galdino, A. S. et al 2011). The multifunctional nature of these peptides makes them appealing for therapeutic use, as targeting multiple cellular pathways in living bacteria is known to reduce the selective pressure for resistance development.

In this study, six peptides that are preferentially cationic, have a-helix, and have more than 30% hydrophobic residues been developed and synthesized based on the features of natural AMPs. One of them, SK1260, demonstrated robust immunomodulatory activity in innate immune cells as well as a potent inhibitory effect against bacteria without hemolyzing human blood red cells. Peptide SK1260 could be a promising candidate for the treatment of infectious diseases.

## 2. Materials and Methods

### 2.1. Design of Peptide

The antibacterial peptides were designed using the APD (http://aps.unmc.edu/AP/main.php) [Z. Wang et al. 2004 and G. Wang et al. 2009] and predicted using (AxPEP server: https://cbbio.online/AxPEP/). The peptide sequences were chosen based on the following rules: (1) containing positive charged amino acids, (2) containing *α*-helix, and (3) containing hydrophobic amino acids. The sequences of six designed peptides are shown in Table 1.

**Table 1:**
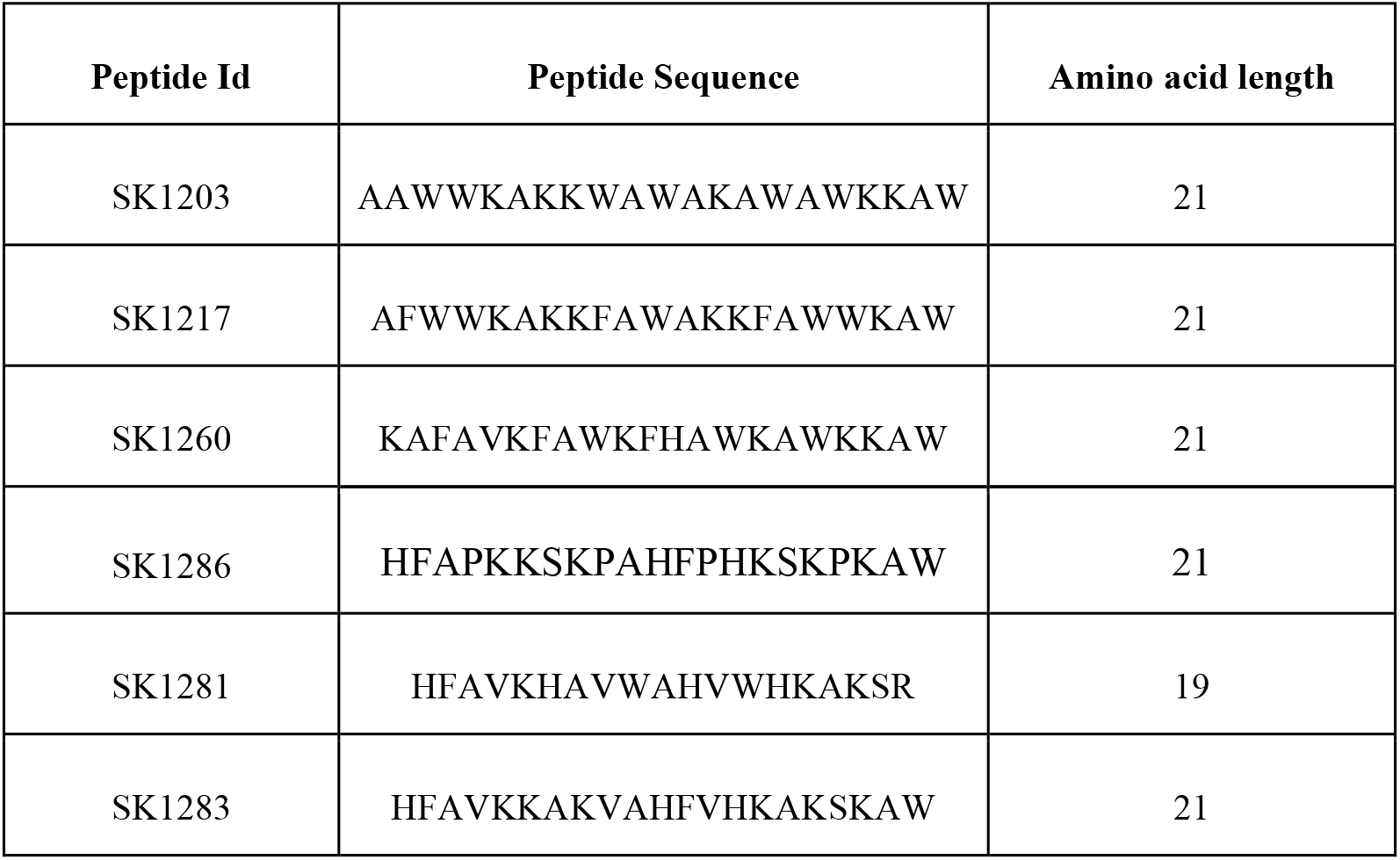
The sequence of the designed peptides

### 2.2. The Strains and Growth Conditions

One Gram-positive bacteria, and 3 Gram-negative bacteria, were selected to measure the antibacterial activity of peptides. The reference strains were provided by the Department of Microbiology, Kakatiya medical college, Warangal, and were stored at 4∘C until use. The species and culture conditions of these bacterial strains are summarized in Table 2.

**Table 2:** Bacterial strains and culture conditions

| Bacterial Strain | Gram Stain | Species | Culture Conditions |
| --- | --- | --- | --- |
| E. coli ATCC 8739 | Gram-negative | Escherichia coli | Growth in LB broth at 37°C |
| S. aureus ATCC 6538 | Gram-positive | Staphylococcus aureus | Growth in nutrient agar at 37°C |
| K. pneumoniae ATCC 700603 | Gram-negative | Klebsiella pneumoniae | Growth in nutrient broth at 37°C |
| P. aeruginosa ATCC 9027 | Gram-negative | Pseudomonas aeruginosa | Growth in TSB medium at 37°C |

### 2.3. Peptide Synthesis and Purification

Peptides were synthesized by the Fmoc (N-[9-fluorenyl]-methoxycarbonyl) chemistry according to the literature procedure [G. B. Fields et al. 1999 and L. P. Miranda et al. 1990]. Protected amino acids were coupled by in situ activation with N,N-diisopropylethylamine (DIEA), and N-hydroxy benzotriazole (HOBt). Deprotection was performed with 20% piperidine in N,N-dimethylformamide (DMF). The protected side chains of amino acid residues and the cleavage of the peptide from the solid support were performed by 95% trifluoroacetic acid (TFA)/2.5% triisopropylsilane (TIS)/2.5% water for 1 h at room temperature. After cleavage from resin, the peptides were purified by preparative reverse-phase HPLC (BioLogic Duoflow system) on a Kromasil C18 column (250 × 10 mm, particle size 5 *μ*m, pore size 100 A °). The elution was achieved with a linear gradient of 0.05% TFA in 5% methanol (A) and 0.05% TFA in 95% methanol (B) at a flow rate of 5 mL/min (57–64% B in 30 min). The main peak was pooled, lyophilized, and stored at −20∘C. The purity of the peptide was evaluated using analytical reverse-phase HPLC (Shimadzu LC-10AT) on a Lichrospher C18 column (250 × 4.6 mm) in the same mobile phase with a linear gradient at a flow rate of 1 mL/min (49–54% B in 20 min). The synthetic peptides were confirmed by electrospray mass spectrometry.

### 2.4. Measurement of Antibacterial Activity

The minimum inhibitory concentrations (MICs) of the AMPs were measured in 96-well microtiter plates according to the Clinical and Laboratory Standards Institute (CLSI) [Wikler MA et al. 2009]. Briefly, liquid Mueller-Hinton-II medium containing various concentrations of AMPs (100, 50, 25, 12.5, 6.25, 3.125, 1.56, and 0.75 *μ*g/mL) is inoculated with a defined number of cells (approx. 10^5^ CFU/ml) in 96-well microtiter plates (polypropylene), whereas each plate also includes a positive growth control and a negative control (sterile control). Ciprofloxacin at a concentration range of 0.5 to 1.0 μg/mL was used as the reference antibiotic. After incubation, the MIC is determined by the lowest concentration showing no visible growth. All MICs were determined in two independent experiments performed in duplicate.

### 2.5. Measurement of Hemolytic Activity (MHC)

The hemolytic activity of the peptides was assayed against human red blood cells (hRBCs) by measuring the amount of hemoglobin released after treatment. The hRBCs, freshly collected from a healthy volunteer in polycarbonate tubes containing heparin, were washed three times in sterile phosphate-buffered saline (PBS) and centrifuged at 2,000 × g for 5 min or until the supernatant became clear. The hRBCs were diluted to a final concentration of 2% (vol/vol), then 50 µl of the hRBCs suspension was incubated with 50 µl of different concentrations (0.98 to 250 μg/ml) of a peptide dissolved in PBS. After 1 h of incubation at 37 °C, intact hRBCs were pelleted by centrifugation at 2,000 × g for 10 min. The supernatant was transferred to a new 96-well plate and the release of hemoglobin was monitored by measurement of absorbance at 405 nm using a Multiskan FC microplate reader. The hRBCs in PBS only (ODBlank) and in 0.1% Triton X-100 (ODTriton X-100) were employed as negative (0% hemolysis) and positive (100% hemolysis) controls, respectively. The percentage of hemolysis was calculated according to the following equation:

%Hemolysis=(ODSample−ABlank)/(ODTritonX−100−ODBlank)×100

### 2.6. Animals, Cell Lines, and Reagents

Male C57BL/6J mice (6-8 weeks) were obtained from the Animal Facility of Jeeva Life Sciences, Hyderabad. The original breeding colonies were obtained from Jackson Labs, USA. The animals were maintained in a pathogen-free condition. All the procedures for animal experiments were approved by the Institutional Animal Ethics Committee (IAEC) and performed in accordance with the Committee for Control and Supervision of Experiments on Animals (CCSEA) guidelines. Mouse, bovine, and human macrophage cell lines RAW264.7, BoMac, and THP-1, respectively, were initially purchased from the American Type Culture Collection (Manassas, VA, United States). Cells were cultured in a specified medium [Dulbecco’s Modified Eagle Medium (DMEM) or Roswell Park Memorial Institute (RPMI) 1640, Sigma, United States] supplemented with 10% Fetal bovine serum (FBS) (Invitrogen, Carlsbad, CA, United States), penicillin (100 U/ml), and streptomycin (100 mg/ml) and maintained at 37°C, 5% CO2. Cells were stimulated by adding 500 ng/mL LPS (E. coli 0111:B4) (Sigma, USA) into the medium specified above. Peptides were added at two different concentrations (2 and 4µg/mL) 30 min after the addition of LPS. After 24Hrs of incubation, plates were centrifuged for 6 min at 400× g and the supernatants were collected and kept frozen at −20 °C until used for analysis of IL-10, IL-6 and TNF-α production by ELISA. Interleukin 6 (IL-6), Interleukin 10 (IL-10), and tumor necrosis factor–α (TNF-α) cytokines sandwich ELISA kits specifically for human or mouse or bovine were purchased from the R&D system.

### 2.7. Cytotoxicity assays

Cellular cytotoxicity was measured by a colorimetric assay that makes use of thiazolyl blue tetrazolium bromide (MTT: Sigma, USA). Sub-confluent monolayer culture of RAW264.7, BoMac, and THP-1 cells was collected by scraper in supplemented DMEM (Invitrogen, USA). Cells were seeded in 96-well microtiter plates at a density of 1.0 × 10^5^ cells per well, with different concentrations of peptide (1 to 10 µg/mL). Cells were incubated at 37 °C in the presence of 5% CO_2_ for 48 h. Following incubation, MTT was added to the cells (10 μL at 5 mg.mL−1). The plate was incubated for 4 hours in the presence of 5% CO2 at 37 °C. Formazan crystals were dissolved by the addition of 100 μL of 100% DMSO (Mallinckrodt Chemical, USA) per well. Plates were then gently swirled for 5 min at room temperature to dissolve the precipitate. Absorbance was monitored at 575 nm using a microplate spectrophotometer (Victor X, PerkinElmer, Germany). Maximum cytotoxicity (100%) was determined by cells incubated with 1% Triton X-100, PBS was used as a negative control.

### 2.8. ELISA

Cultured cells or peritoneal lavage samples were centrifuged at 1000 × g for 10 min to obtain cell-free samples and stored at −20 °C. Cytokine levels were measured by ELISA using anti-mouse IL-6, IL-10, GM-CSF, MCP-1, INF-γ, and TNF-α (R&D Systems). Cytokine levels were analysed according to the manufacturer’s instructions.

### 2.9 Protein-Protein docking studies

Amino acid sequences of 6 peptides (SK1203, 1217, 1260, 1281,1283, 1286) were submitted to blastp to find a structural homologue as a template used for homology modeling. However, no significant structural homology was observed. Hence, the amino acid sequences of all 6 peptides were submitted for 3-dimensional structural prediction using the I-TASSER server (Roy A et al. 2010) which generated the five top best-predicted structures with high C-score. A structure in agreement with the Ramachandran plot was selected for further energy minimization using the steepest descent algorithm in NOMAD-REF online program (http://lorentz.dynstr.pasteur.fr/docking/submission.php)(Lindahl, E et al. 2006). The energy-minimized structures were used to create a docking model on human or bovine or mice TLR4 structures using the ZDOCK server (http://zdock.umassmed.edu)(Pierce, B. G. et al. 2014). The atomic coordinates of human, bovine, and mice TLR4 were taken from the crystal structure of hTLR4 complex (3FXI), bTLR4 (3RG1), and mTLR4 (3VQ1) respectively as static structures (receptor) for docking. The docking program generated the top 10 scoring complexes based on shape complementarity, electrostatics, and rigid body.

### 2.10 Mouse systemic infection model

Male C57BL/6 mice, aged 6 weeks and matched by weight, were used for the study. Log phase bacterial inoculum of either E. coli ATCC 8739 or S. aureus ATCC 6538 was prepared and washed before resuspending in 0.1 ml of sterile PBS. The bacterial suspension was then injected intraperitoneally (i.p.) into the mice at a dose of 5.0 × 10^8^ CFU/mice. After 3 hours of bacterial infection, the mice (n=8) were treated with either peptide SK1260 (2 or 4 mg/kg) by i.p. administration or received an equivalent volume of PBS as the control group. At 4- and 24-hours post-treatment, peritoneal lavage was performed by washing the peritoneal cavity with 5 ml of sterile PBS. The lavage fluid obtained was used for further analysis. The number of leukocytes in the lavage fluid was counted by diluting appropriately and using a automated cell counter. Cytokine levels in the lavage fluid, including GM-CSF, IFN-γ, and MCP-1, were measured using cytokine-specific assays ELISA as per manufacturer’s instructions.

### 2.11 Statistical Analysis

For all the experiments, wherever required, Student’s t-test and one way ANOVA were executed for the analysis of the results. The data were represented as the mean of triplicates ± SEM. p < 0.05 was considered as significant. GraphPad Prism software v9.1.0 (GraphPad Software, USA) was used for all statistical analyses

## 3. Results and Discussion

### 3.1. Peptide Design and AMP Prediction

Six peptides were designed based on characteristics of natural AMPs, including small size, cationicity, amphipathicity, and *α*-helix. Five out of the six designed peptides are classified as antimicrobial peptides (AMPs) by the AxPEP server (Pratiti Bhadra et al. 2018). The peptide with the Peptide Id “SK1281” is predicted to be a Non-Antimicrobial Peptide (NAMP) and the physicochemical properties of the peptides, such as molecular weight, cationicity, hydrophobicity, protein-binding potential, and GRAVY score, provide insights into their potential functions and properties Table 3, these predictions are based on the sequence-based classification method employed by the AxPEP server, and further experimental validation may be necessary to confirm the antimicrobial activity of these peptides.

**Table 3:**
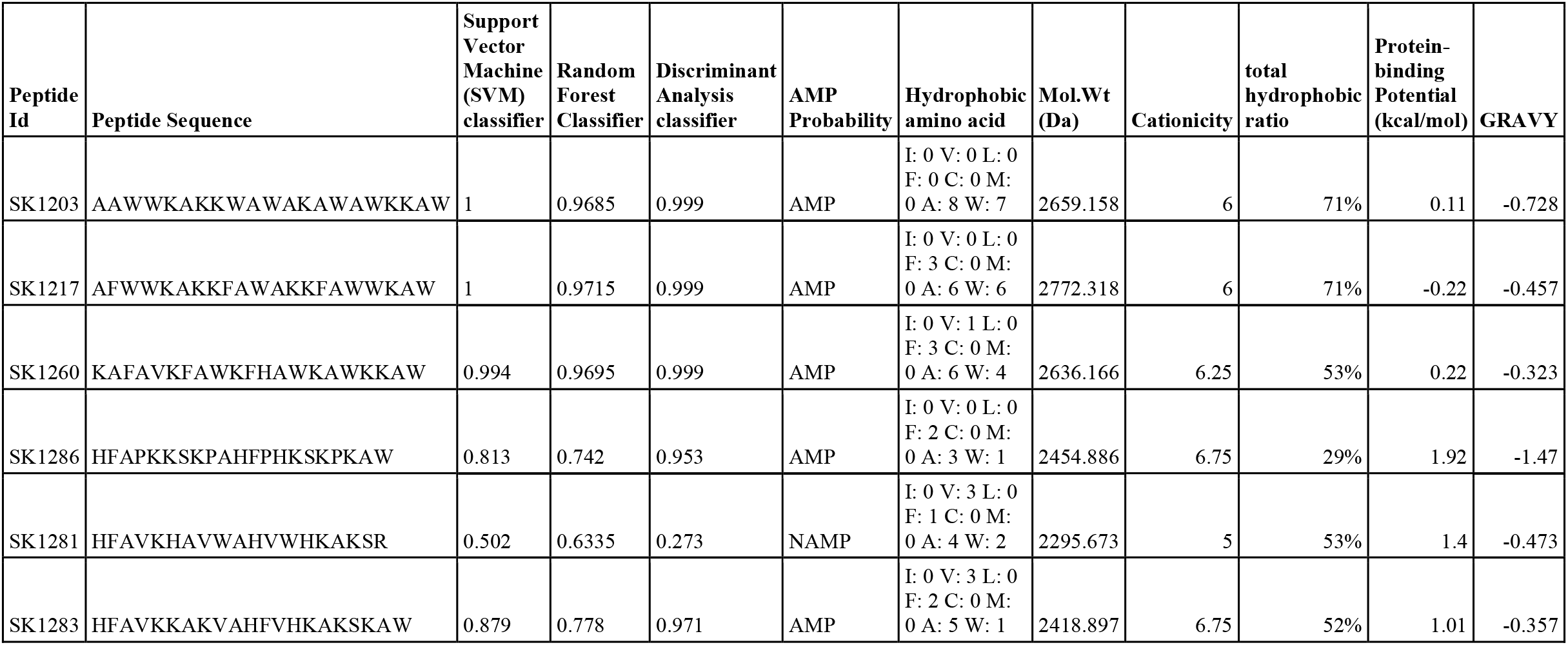
Physicochemical properties of designed peptides

### 3.2. Hemolytic Activity of Designed Peptides on Human Red Blood Cells (RBCs)

The hemolytic activities of six peptides (SK1203, SK1217, SK1260, SK1281, SK1283, and SK1286) were evaluated using a hemolytic assay with human blood red cells (RBCs) at various concentrations. The peptides were tested at concentrations ranging from 0 μg/mL to 100 μg/mL. The percentage of hemolysis observed at each peptide concentration was recorded. Among the designed peptides, SK1203, SK1217, SK1260, and SK1281 displayed minimal hemolytic activity across all tested concentrations, suggesting a low propensity to cause RBC lysis. Whereas, SK1286 and SK1283 exhibited significant hemolytic effects at higher concentrations. At 100 μg/mL, SK1286 caused complete hemolysis (102%), indicating a potent cytotoxic effect on RBCs. SK1283 also demonstrated concentration-dependent hemolysis, reaching 39% at 3.13 μg/mL and 102% at 100 μg/mL. Melittin, a well-known hemolytic peptide, exhibited dose-dependent hemolytic activity, with an increasing percentage of hemolysis as the peptide concentration increased (Figure 1).

**Figure 1:**
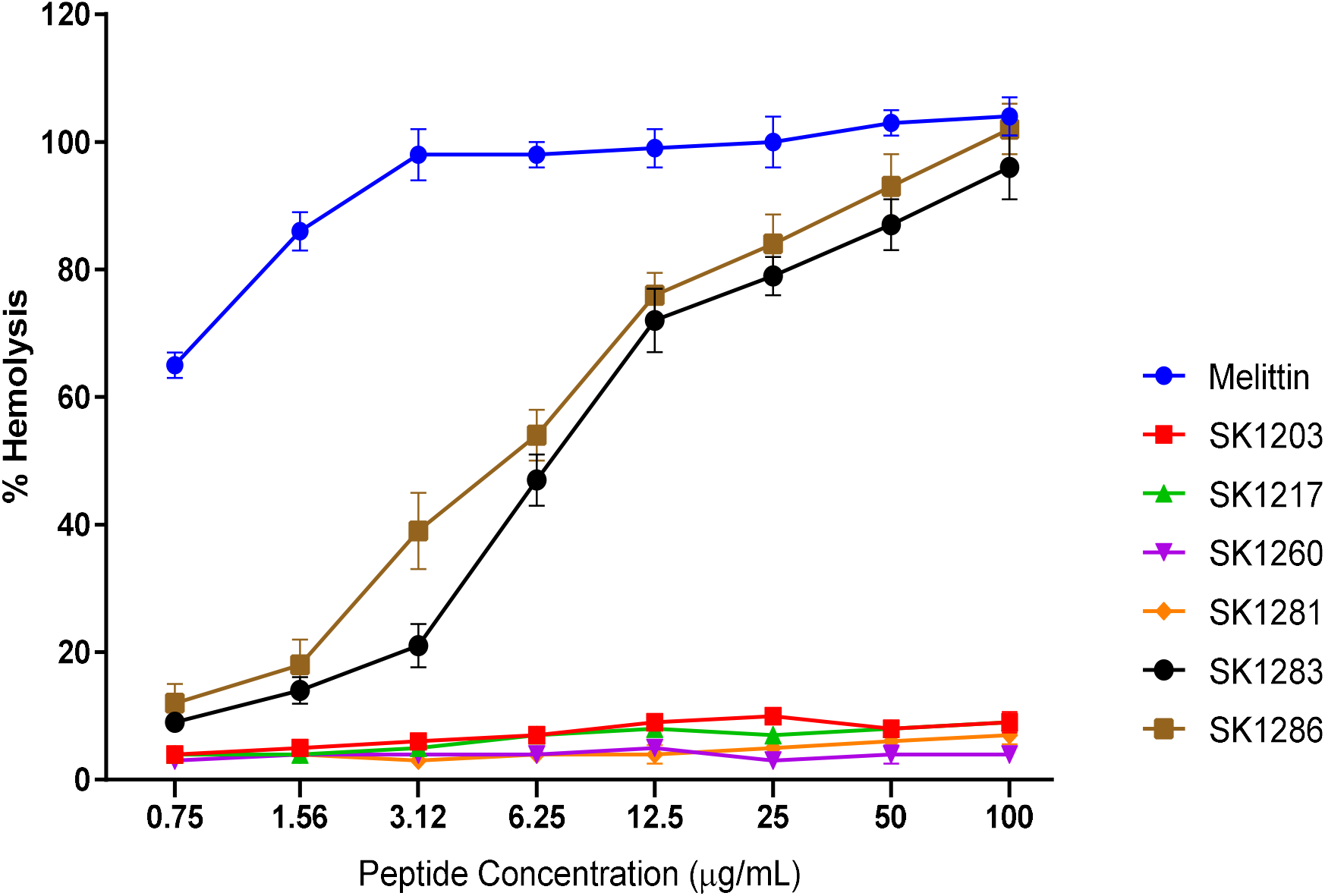
Hemolytic Activity of Designed Peptides. The hemolytic assay was performed using human blood red cells (RBCs) at various concentrations of the peptides, ranging from 0 μg/mL to 100 μg/mL. The percentage of hemolysis observed at each peptide concentration is depicted.

### 3.3. Minimum Inhibitory Concentrations (MICs) of Designed Peptides against Different Microorganisms

Minimum Inhibitory Concentrations (MICs) of six peptides (SK1203, SK1217, SK1260, SK1281, SK1283, SK1286) against different microorganisms are summarized in Table 5. SK1203 displayed moderate antimicrobial activity, with MIC values of 12.5 ug/mL against Escherichia coli (E. coli) and 25 ug/mL against both Staphylococcus aureus (S. aureus) and Pseudomonas aeruginosa (P. aeruginosa). SK1217 exhibited stronger antimicrobial activity, with MIC values of 6.25 ug/mL against E. coli and Klebsiella pneumoniae (K. pneumoniae), 3.125 ug/mL against S. aureus, and 12.5 ug/mL against P. aeruginosa. SK1260 demonstrated potent antimicrobial activity, with MIC values of 3.13 ug/mL against E. coli, S. aureus, K. pneumoniae, and P. aeruginosa. SK1283 showed moderate activity, with MIC values of 12.5 ug/mL against E. coli and P. aeruginosa. SK1286 exhibited moderate to potent antimicrobial activity, with MIC values of 12.5 ug/mL against E. coli and P. aeruginosa, and 6.25 ug/mL against S. aureus, and no significant activity against K. pneumoniae, Table 4. Overall, these findings indicate that the designed peptides possess varying degrees of antimicrobial activity against the tested microorganisms. SK1217 and SK1260 demonstrate stronger antimicrobial potential, while SK1203, SK1283, and SK1286 exhibit moderate activity.

**Table 4:** Minimum Inhibitory concentrations of the designed peptides

|  | MIC (ug/mL) |  |  |  |  |  |
| --- | --- | --- | --- | --- | --- | --- |
|  | SK1203 | SK1217 | SK1260 | SK1281 | SK1283 | SK1286 |
| **E.coli** | 12.5 | 6.25 | 3.13 | No activity | 12.5 | 12.5 |
| **S.aureus** | 25 | 3.125 | 3.13 | No activity | 12.5 | 6.25 |
| **K.pneumonia** | No activity | 6.25 | 3.13 | No activity | No activity | No activity |
| **P.aeruginosa** | 25 | 12.5 | 3.13 | No activity | 12.5 | 6.25 |

**Table 5:**
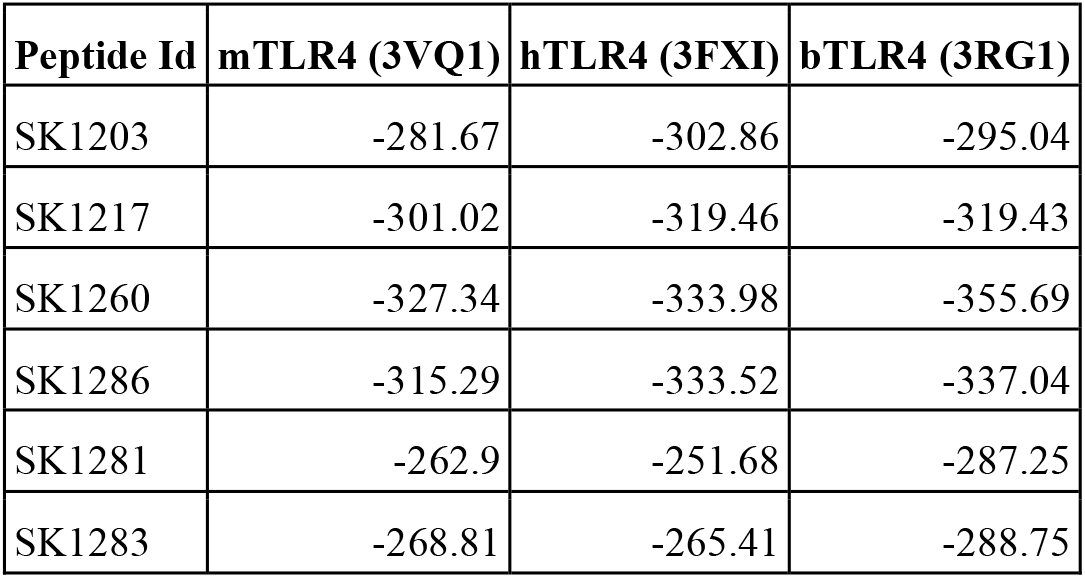
Molecular docking

| Peptide Id | mTLR4 (3VQ1) | hTLR4 (3FXI) | bTLR4 (3RG1) |
| --- | --- | --- | --- |
| SK1203 | -281.67 | -302.86 | -295.04 |
| SK1217 | -301.02 | -319.46 | -319.43 |
| SK1260 | -327.34 | -333.98 | -355.69 |
| SK1286 | -315.29 | -333.52 | -337.04 |
| SK1281 | -262.9 | -251.68 | -287.25 |
| SK1283 | -268.81 | -265.41 | -288.75 |

**Fig 2**
| Peptide Id | mTLR4 (3VQ1) | hTLR4 (3FXI) | bTLR4 (3RG1) |
| --- | --- | --- | --- |
| SK1203 | -281.67 | -302.86 | -295.04 |
| SK1217 | -301.02 | -319.46 | -319.43 |
| SK1260 | -327.34 | -333.98 | -355.69 |
| SK1286 | -315.29 | -333.52 | -337.04 |
| SK1281 | -262.9 | -251.68 | -287.25 |
| SK1283 | -268.81 | -265.41 | -288.75 |

### 3.4 Molecular Docking Analysis of Designed Peptides with TLR4/MD-2 Receptor Proteins

Molecular docking was employed to investigate the binding patterns and affinity of the designed peptides (SK1203, SK1217, SK1260, SK1286, SK1281, and SK1283) with the receptor proteins mouse TLR4/MD-2 (3VQ1), human TLR4/MD-2 (3FXI), and bovine TLR4/MD-2 (3RG1). The docking scores, representing the predicted affinity between each peptide and the respective receptor protein, are summarized in Table 5. Lower docking scores indicate stronger binding affinity between the peptide and the receptor protein. Based on the docking scores, SK1260 demonstrated the highest affinity for all three receptors, with docking scores of −327.34, −333.98, and −355.69 for mTLR4/MD-2 (3VQ1), hTLR4/MD-2 (3FXI), and bTLR4/MD-2 (3RG1), respectively. SK1217 and SK1286 also displayed favourable binding affinities across the receptors Figure 2. These findings suggest that SK1260, SK1217, and SK1286 have promising potential immunomodulatory peptides candidates.

**Figure 2:**
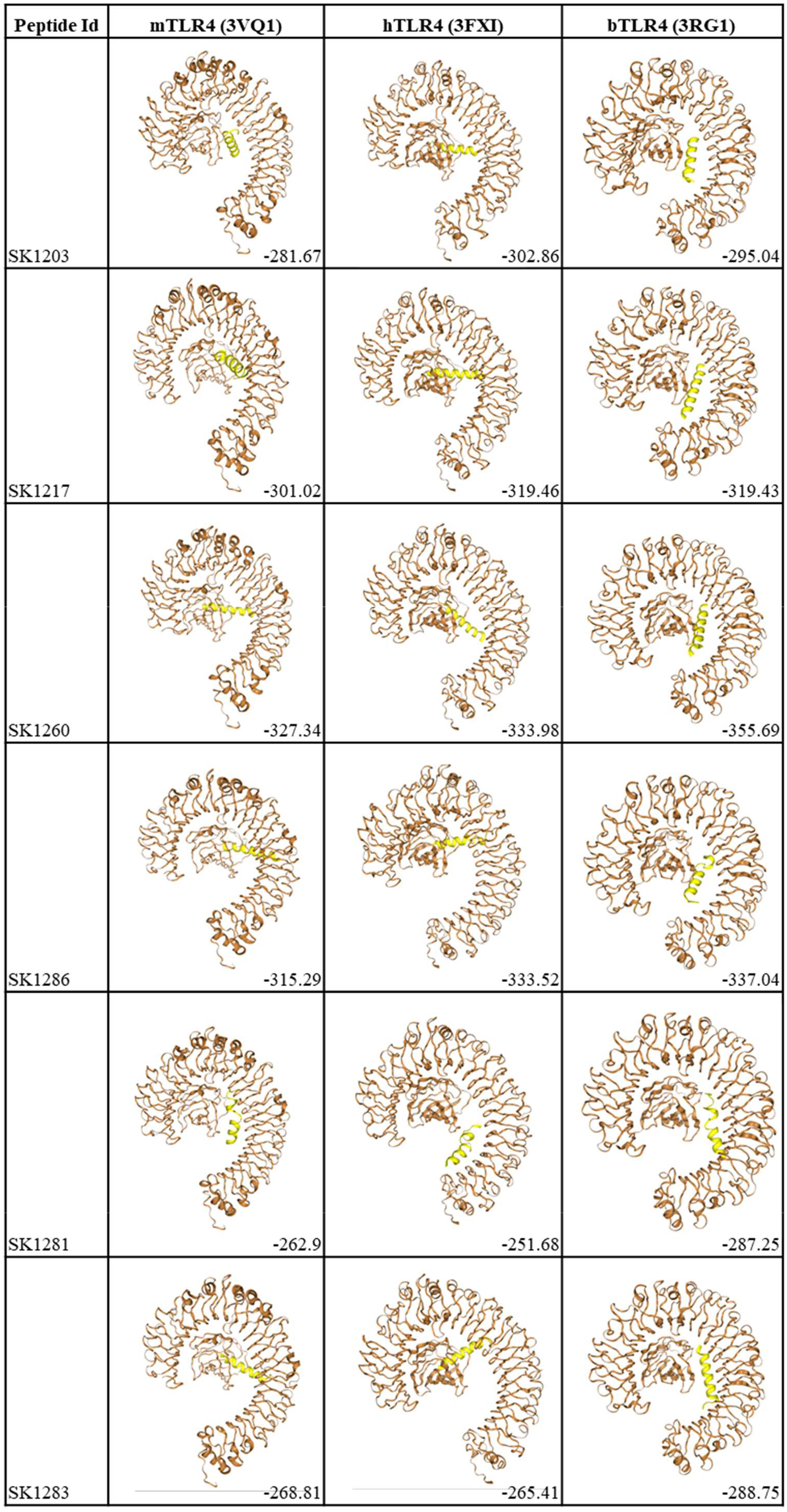
Binding Affinity of Designed Peptides with TLR4/MD-2 Receptors. Molecular docking analysis, examining the binding patterns and affinity of six designed peptides (SK1203, SK1217, SK1260, SK1286, SK1281, and SK1283) with three receptor proteins: mouse TLR4/MD-2 (3VQ1), human TLR4/MD-2 (3FXI), and bovine TLR4/MD-2 (3RG1).

### 3.5 Modulation of Innate Immune Response by Cationic Antimicrobial Peptides in Macrophage Cell Lines

The effects of cationic antimicrobial peptides (AMPs) SK1203, SK1217, SK1260, and SK1281 on the modulation of the innate immune response were examined in three different cell lines: murine macrophage-like RAW 264.7 cells, human macrophage cell line (THP-1), and bovine macrophage cell line (BoMac). The cells were stimulated with 500 ng/mL lipopolysaccharide (LPS) and treated with various concentrations of the AMPs. The production of pro-inflammatory cytokines (IL-6 and TNF-α) and anti-inflammatory cytokine (IL-10) was measured. In all three cell lines, LPS stimulation led to a significant increase in IL-6 and TNF-α levels compared to the control group (PBS treatment). However, treatment with the AMPs resulted in a concentration-dependent reduction in IL-6 and TNF-α production while maintaining stable IL-10 levels (Figure 3A-C). Among the peptides tested, SK1260 showed the most potent inhibitory effect on IL-6 production in RAW 264.7 and THP-1 cells, while SK1217 exhibited the strongest inhibitory effect on TNF-α production in BoMac cells. These findings suggest that these cationic AMPs have the potential to modulate the innate immune response by suppressing pro-inflammatory cytokines while preserving anti-inflammatory cytokine production in different cell types.

**Figure 3.**
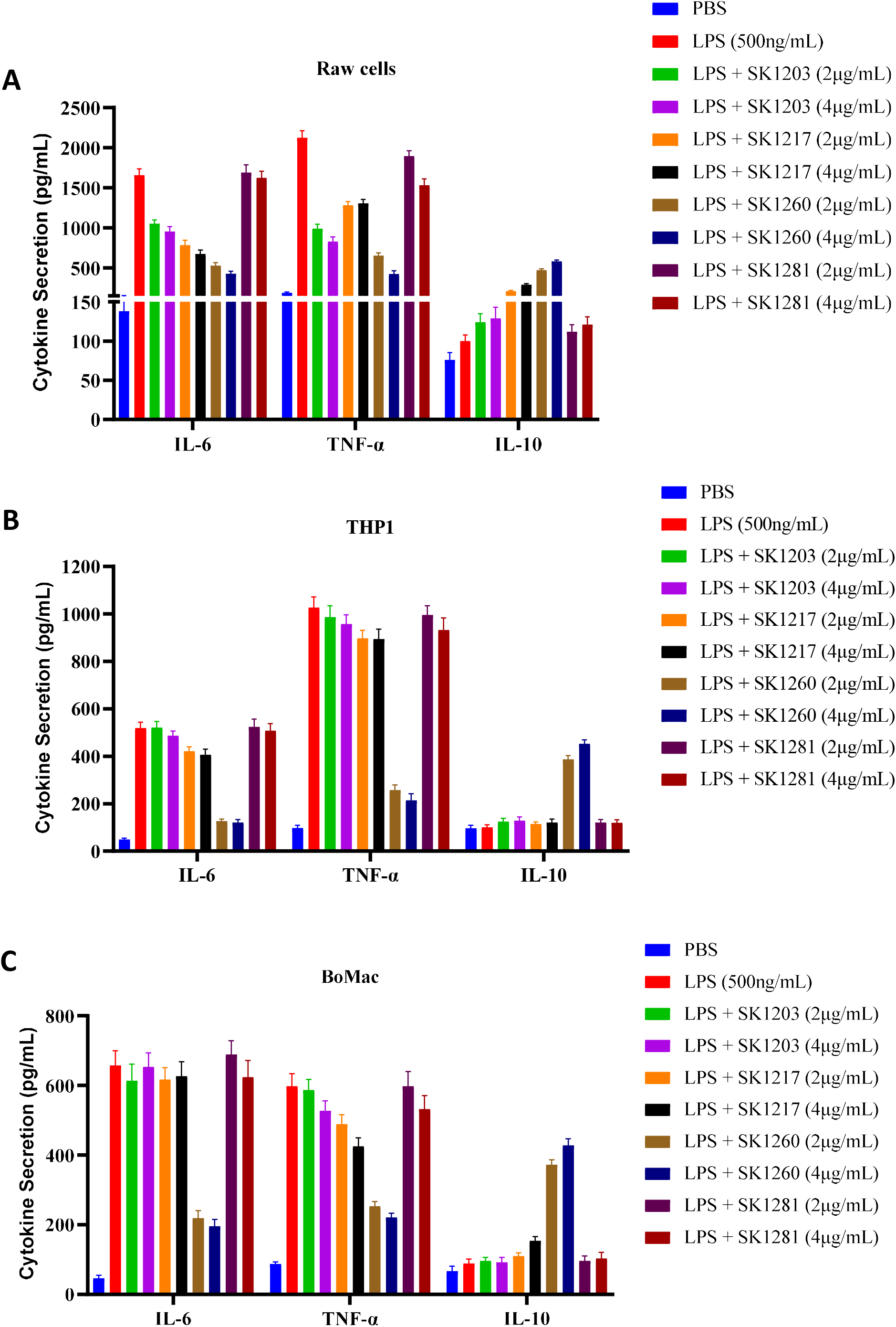
Depicts the effects of cationic antimicrobial peptides (AMPs) SK1203, SK1217, SK1260, and SK1281 on the modulation of the innate immune response in different cell lines. (A) RAW 264.7 cells, (B) THP-1 cells, and (C) BoMac cells were stimulated with 500 ng/mL lipopolysaccharide (LPS) and treated with various concentrations of the AMPs. The production of pro-inflammatory cytokines IL-6 and TNF-a, as well as the anti-inflammatory cytokine IL-10, was measured.

### 3.6 Cell Viability Assay of SK1260 on RAW264.7, THP-1, and BoMac Cells

After determining the modulation of the innate immune response by peptide SK1260, we further investigated its cytotoxic effects on mouse macrophage cells RAW264.7, human macrophage cell line (THP-1), and bovine macrophage cell line (BoMac). Cell viability was assessed using the MTT assay after 24 hours of treatment with different concentrations of SK1260. The results showed that SK1260 exhibited minimal cytotoxicity at all tested concentrations. In RAW264.7 cells, cell viability remained high, with 99.8% at 1 μg/mL, 97.8% at 2 μg/mL, 98.9% at 5 μg/mL, and 96.8% at 10 μg/mL of SK1260. Similarly, in THP-1 cells, viability was well-preserved, with 98.2% at 1 μg/mL, 97.6% at 2 μg/mL, 94.8% at 5 μg/mL, and 96.2% at 10 μg/mL of SK1260. BoMac cells also exhibited high viability, with 99.3% at 1 μg/mL, 96.2% at 2 μg/mL, 97.2% at 5 μg/mL, and 96.7% at 10 μg/mL of SK1260 (Figure 4). These results demonstrate that SK1260 has minimal cytotoxic effects on all three cell lines, suggesting its potential as a safe and biocompatible peptide for further therapeutic applications

**Figure 4:**
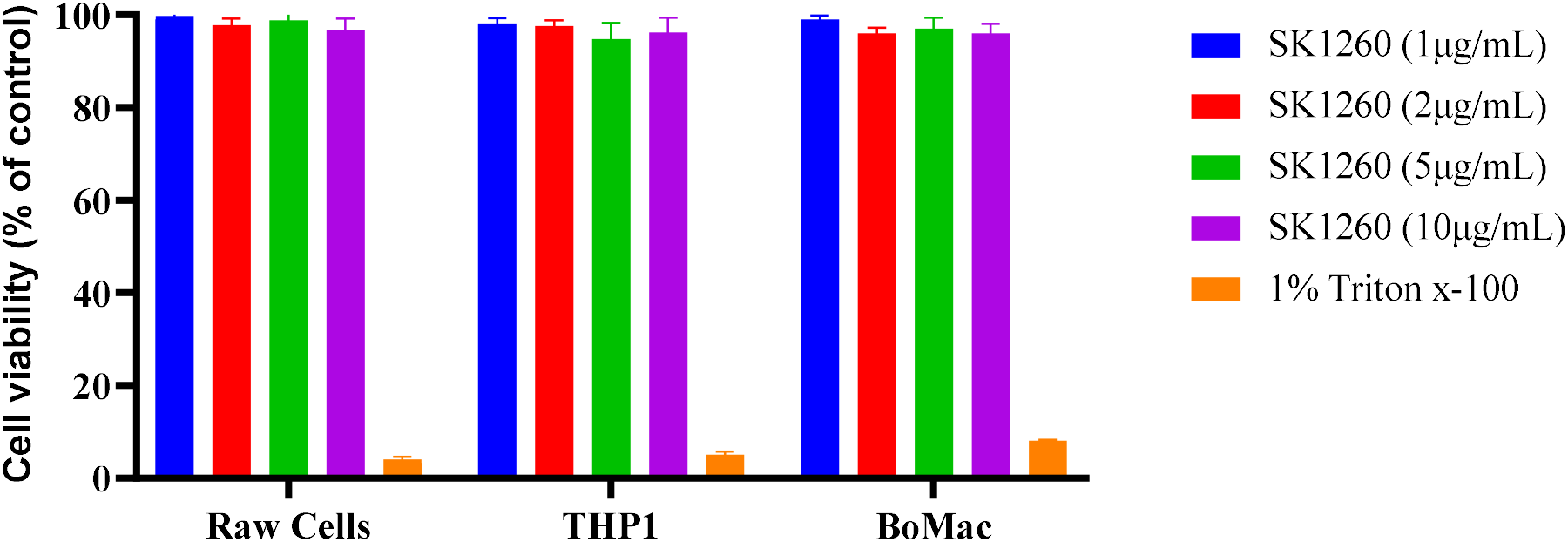
Cell viability assay of peptide SK1260 on (A) RAW264.7 cells, (B) THP-1 cells, and (C) BoMac cells. The MTT assay was performed after 24 hours of treatment with different concentrations of SK1260. The results demonstrate that SK1260 exhibited minimal cytotoxicity on all tested cell lines.

### 3.7 SK1260 mediated immune modulation in mice

Next, we investigated the leukocyte count in the peritoneal lavage fluid of male C57BL/6 mice infected with either E. coli or S. aureus and treated with peptide SK1260. At 4 hours post-treatment, the control group (PBS) showed a mean leukocyte count of 0.8 × 10^6^ (± 0.2) in the lavage fluid. In the E. coli-infected group treated with PBS, the mean leukocyte count was 0.7 × 10^6^ (± 0.12). However, treatment with SK1260 at a concentration of 2mg/mL resulted in a significant increase in leukocyte count, with a mean of 1.2 × 10^6^ (± 0.12). Similarly, in the S. aureus-infected group, treatment with SK1260 at 2mg/mL led to a higher leukocyte count (mean = 1.4 × 10^6^, ± 0.12) compared to the group treated with PBS (mean = 0.7 × 10^6^, ± 0.18), Figure 5A&B. Furthermore, the cytokine levels (GM-CSF, INF-g, and MCP-1) in the peritoneal lavage fluid were measured. At 4 hours post-treatment, the control group (PBS) exhibited basal levels of cytokines. In the E. coli-infected group treated with SK1260 (2mg/mL), there was an elevation in GM-CSF (mean = 627, ± 38), INF-g (mean = 534, ± 35), and MCP-1 (mean = 728, ± 43) compared to the PBS-treated group (GM-CSF mean = 324, INF-g mean = 128, MCP-1 mean = 126). Similarly, in the S. aureus-infected group treated with SK1260 (2mg/mL), higher levels of GM-CSF (mean = 572, ± 41), INF-g (mean = 434, ± 41), and MCP-1 (mean = 628, ± 51) were observed compared to the PBS-treated group (GM-CSF mean = 243, INF-g mean = 218, MCP-1 mean = 216), Figure 6A&B. These results demonstrate that SK1260 treatment can modulate leukocyte recruitment and cytokine production in response to bacterial infection, indicating its potential as a therapeutic agent for immune modulation in infectious diseases.

**Figure 5:**
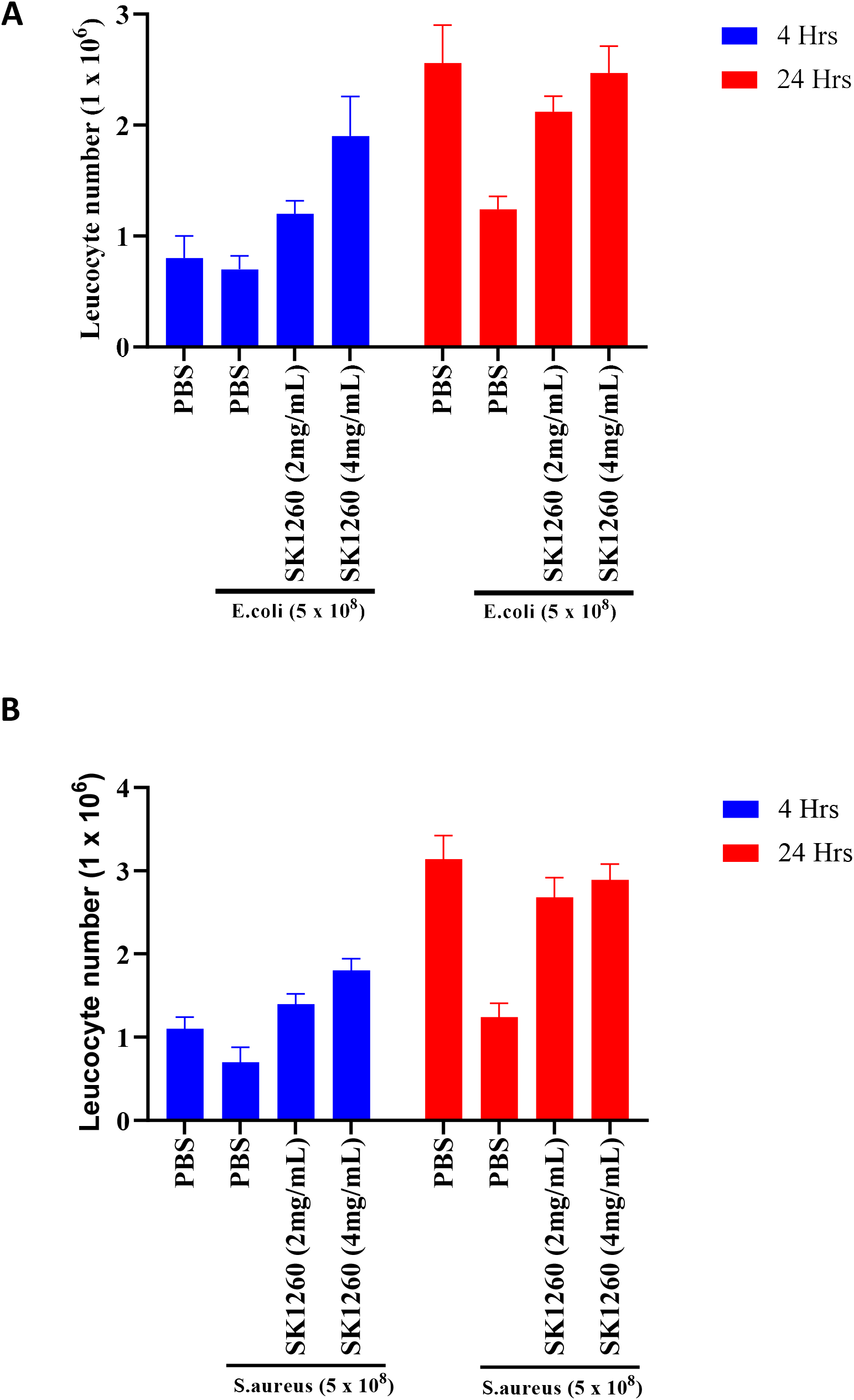
Leukocyte counts in the peritoneal lavage fluid of mice infected with E. coli (a) or S. aureus (b) and treated with different concentrations of SK1260 or PBS.

**Figure 6:**
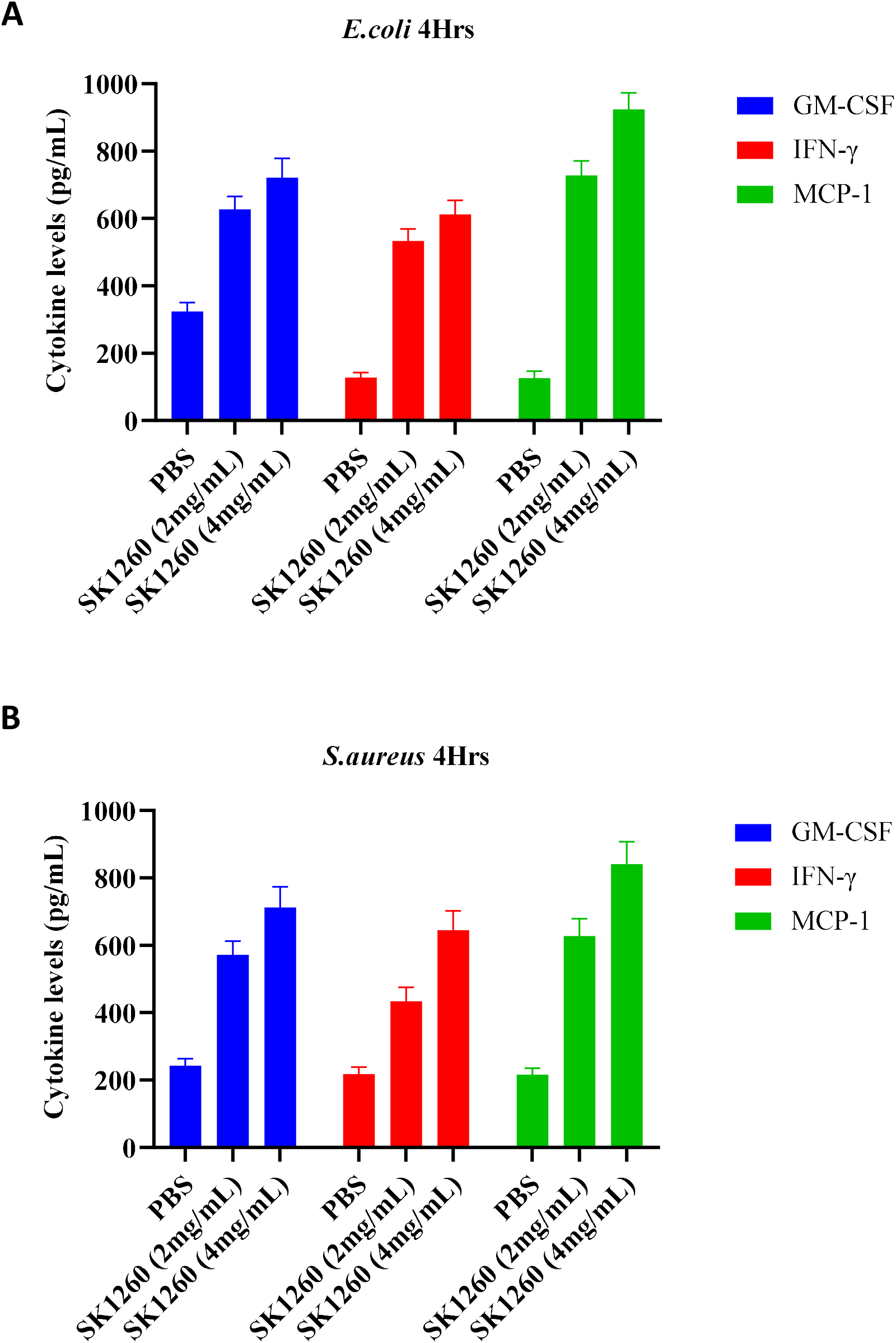
Cytokine levels (GM-CSF, IFN-γ, and MCP-1) in the peritoneal lavage fluid of mice infected with E. coli (a) or S. aureus (b) and treated with different concentrations of SK1260 or PBS.

## 4. Discussion

Our study focused on the design, characterization, and evaluation of six peptides with potential antimicrobial and immunomodulatory properties. The design of these peptides was inspired by the characteristics of natural antimicrobial peptides (AMPs) that are known to exhibit dual antimicrobial and immunomodulatory activities. AMPs have garnered significant attention in recent years due to their ability to not only directly kill or inhibit the growth of microorganisms but also modulate the immune response (Hancock, R. E. W. et al., 2016). By incorporating features such as small size, cationicity, amphipathicity, and α-helical structure, we aimed to create peptides with similar functional properties.

Several AMPs with dual antimicrobial and immunomodulatory activities have been identified and studied. For instance, LL-37, derived from human cathelicidin, exhibits antimicrobial activity against a broad range of pathogens, including bacteria, fungi, and viruses (Wang, G., 2014). LL-37 also displays immunomodulatory effects by promoting wound healing, enhancing chemotaxis of immune cells, and stimulating the production of cytokines and chemokines (Gallo, R. L. et al., 2012). Similarly, human β-defensins, such as HBD-2 and HBD-3, possess both antimicrobial and immunomodulatory functions (Yang, D. et al., 2002). These peptides not only exert direct antimicrobial effects but also regulate the production of pro-inflammatory cytokines, influence antigen presentation, and contribute to immune cell recruitment (Niyonsaba, F. et al., 2009).

The dual functionality of AMPs provides several advantages in combating infections and promoting host defense. Firstly, their antimicrobial activity allows for direct killing or inhibition of pathogenic microorganisms, reducing the risk of infection (Hancock, R. E. W. et al., 2016). This antimicrobial efficacy is particularly important considering the rise in antibiotic resistance among pathogens. Secondly, the immunomodulatory effects of AMPs contribute to shaping the immune response, enhancing host defense mechanisms, and promoting resolution of inflammation (Gallo, R. L. et al., 2012). By modulating the production of cytokines and chemokines, AMPs can fine-tune the immune response, preventing excessive inflammation or immunopathology. Moreover, AMPs can influence immune cell functions, such as chemotaxis, phagocytosis, and antigen presentation, thereby enhancing the overall immune defense against infections (Hancock, R. E. W. et al., 2016).

Despite the advantages of AMPs with dual antimicrobial and immunomodulatory activities, there are certain limitations and challenges associated with their use. One of the main challenges is their susceptibility to degradation by proteases in biological fluids and tissues, which may limit their stability and efficacy (Zasloff, M., 2002). Strategies such as peptide modifications, nanoparticle encapsulation, or delivery systems are being explored to enhance their stability and prolong their half-life (Wang, G., 2014). Another challenge is the potential for cytotoxicity or hemolytic activity, which can limit their therapeutic use (Oddo, A. et al., 2017). It is crucial to carefully evaluate the cytotoxic effects of AMPs and optimize their design to minimize adverse effects on host cells.

In our study, five out of the six designed peptides were classified as antimicrobial peptides (AMPs) based on sequence-based classification using the AxPEP server. This classification suggests that these peptides share sequence similarities or features with known AMPs and may possess antimicrobial activity. AMPs are known for their ability to directly kill or inhibit the growth of various microorganisms, including bacteria, fungi, and viruses (Hancock, R. E. W. et al., 2016). The classification of our designed peptides as AMPs provides an initial indication of their potential antimicrobial efficacy. One peptide, SK1281, was predicted to be a Non-Antimicrobial Peptide (NAMP) by the AxPEP server.

To evaluate the antimicrobial activity of the designed peptides, we performed hemolytic assays and determined their Minimum Inhibitory Concentrations (MICs) against different bacterial strains. Hemolytic assays assess the ability of peptides to cause damage to red blood cells (RBCs) and serve as a measure of potential toxicity. Four out of the six peptides (SK1203, SK1217, SK1260, and SK1281) exhibited minimal hemolytic activity across a range of concentrations, suggesting a low propensity to cause RBC lysis. This is a favorable characteristic for potential therapeutic use, as hemolysis can lead to adverse effects in vivo (Oddo, A. et al., 2017). However, two peptides, SK1286 and SK1283, showed significant hemolytic effects at higher concentrations, indicating a higher potential for toxicity. These findings highlight the importance of evaluating the hemolytic activity of AMPs to ensure their safety for therapeutic applications.

MIC determination revealed varying degrees of antimicrobial activity among the designed peptides. SK1217 and SK1260 demonstrated potent antimicrobial activity against all tested bacterial strains, including Gram-negative Escherichia coli (E. coli) and Pseudomonas aeruginosa (P. aeruginosa), as well as Gram-positive Staphylococcus aureus (S. aureus) and Klebsiella pneumoniae (K. pneumoniae). These peptides exhibited MIC values in the low micromolar range, indicating their effectiveness against a broad spectrum of bacteria. The broad-spectrum antimicrobial activity of AMPs is advantageous as it allows for potential use against diverse pathogens, reducing the need for multiple specific treatments (Hancock, R. E. W. et al., 2016).

To gain insights into the binding patterns and affinity of the designed peptides with receptor proteins involved in immune response regulation, we conducted molecular docking studies. Our results showed that SK1260 displayed the highest affinity for the tested receptors, suggesting its potential as an immunomodulatory peptide. SK1217 and SK1286 also exhibited favorable binding affinities. The interaction of AMPs with receptors, such as Toll-like receptor 4 (TLR4)/MD-2, can trigger downstream signaling pathways that modulate immune responses (Vidal-Limon, A. et al., 2022). By binding to these receptors, the peptides may influence immune signaling, leading to the modulation of pro-inflammatory and anti-inflammatory cytokine production (Gokhale, A. S. et al., 2014). This immunomodulatory property of the peptides adds an additional layer of therapeutic potential, as an exaggerated or dysregulated immune response is implicated in various inflammatory conditions (Liu, J. et al., 2016).

The immunomodulatory effects of the designed peptides were further investigated using macrophage cell lines. These cell lines represent key players in the innate immune response and play a crucial role in recognizing and eliminating microbial pathogens (Alon, R. et al., 2021). Our results demonstrated that the AMPs were capable of modulating the immune response by reducing the production of pro-inflammatory cytokines, such as IL-6 and TNF-α, while maintaining stable levels of the anti-inflammatory cytokine IL-10. SK1260 exhibited the most potent inhibitory effect on IL-6 production, while SK1217 showed the strongest inhibitory effect on TNF-α production. These findings suggest that the designed peptides have the potential to selectively suppress pro-inflammatory cytokines without compromising the anti-inflammatory response. This selective modulation of cytokine production can help restore immune homeostasis and mitigate excessive inflammation, which is often associated with detrimental effects (Yang, L. et al., 2021).

The evaluation of cell viability demonstrated that SK1260 exhibited minimal cytotoxicity in the macrophage cell lines tested. This finding is crucial for the potential clinical application of the peptide, as it suggests its safety and biocompatibility with immune cells. The low cytotoxicity indicates that SK1260 may have a favourable therapeutic window, where it can modulate immune responses without causing harm to host cells (Chen, Y. et al., 2020). The safety profile of AMPs is a critical consideration for their use as therapeutics, and the minimal cytotoxicity observed for SK1260 supports its potential in further preclinical investigations.

In the in vivo evaluation of SK1260, we assessed its effects on leukocyte recruitment and cytokine levels in the peritoneal lavage fluid of infected mice. SK1260 treatment resulted in increased leukocyte counts compared to the control group, indicating enhanced immune cell recruitment to the site of infection. This suggests that SK1260 can promote an effective immune response by facilitating immune cell mobilization to combat bacterial pathogens. Furthermore, SK1260 treatment led to higher levels of cytokines, including GM-CSF, IFN-γ, and MCP-1, which are involved in immune cell activation and coordination of immune responses against infections. These findings demonstrate the potential of SK1260 to modulate cytokine production and enhance immune defences against bacterial pathogens in vivo.

## 5. Conclusion

In conclusion, our study focused on the design and characterization of cationic AMPs with dual antimicrobial and immunomodulatory activities. The designed peptides displayed varying degrees of antimicrobial activity against Gram-negative and Gram-positive bacteria, with SK1217 and SK1260 exhibiting potent broad-spectrum antimicrobial efficacy. The peptides demonstrated minimal hemolytic activity, indicating a low potential for toxicity. Molecular docking studies suggested their potential to interact with receptor proteins involved in immune response regulation, highlighting their immunomodulatory properties. The AMPs effectively modulated cytokine production in macrophage cell lines, selectively suppressing pro-inflammatory cytokines while maintaining anti-inflammatory cytokine levels. SK1260 exhibited minimal cytotoxicity and showed promising effects in promoting immune cell recruitment and cytokine production in infected mice. These findings emphasize the potential of cationic AMPs as therapeutic agents with dual antimicrobial and immunomodulatory properties. Further investigations, including in vivo efficacy studies and toxicity assessments, are warranted to fully explore their clinical applications.

## Acknowledgements

The authors would like to thank the Chancellor, CDU, Dr. Chintalpani V. Purushotham Reddy and Vice chancellor, CDU, Prof. Damodar Gurrapu, for providing the necessary infrastructural facility and support for the execution of the above study. Thanks to Dr. N. Uttam Kumar, Jeeva life sciences for help in animal experiments. SK is supported by CDU institutional fellowship and registered for Ph.D. program at CDU, Hyderabad. MK is supported by UGC fellowship and registered for Ph.D. program at RCB, Faridabad.

## Author’s contribution

Sridhar K, MKT and SC conceived the idea and designed the experiments. SK, Sridhar K, MK performed the experiments. SK, Sridhar K, MK, MKT and Sc analysed the data. Sridhar K made all the figures of manuscript and did the statistical analysis. SK and Sridhar K wrote the initial draft, and MKT and SC edited the manuscript. All authors approved the final version of the manuscript.

## References

1. Alon, R., Sportiello, M., Kozlovski, S. et al. Leukocyte trafficking to the lungs and beyond: lessons from influenza for COVID-19. Nat Rev Immunol 21, 49–64 (2021). https://doi.org/10.1038/s41577-020-00470-2

2. Chen, Y., et al., 2020. Antimicrobial peptides: potential application in liver cancer. Front. Oncol. 10, 515.

3. De la Fuente-Núñez, C., Reffuveille, F., Haney, E. F., Straus, S. K. & Hancock, R. E. W. Broad-spectrum anti-biofilm peptide that targets a cellular stress response. PLoS Pathog. 10, e1004152 (2014).

4. G. B. Fields and R. L. Noble, “Solid phase peptide synthesis utilizing 9-fluorenylmethoxycarbonyl amino acids,” International Journal of Peptide and Protein Research, vol. 35, no. 3, pp. 161– 214, 1990.

5. G. Wang, X. Li, and Z. Wang, “APD2: the updated antimicrobial peptide database and its application in peptide design,” Nucleic Acids Research, vol. 37, supplement 1, pp. D933–D937, 2009.

6. Galdino, A. S. et al. Biochemical and Structural Characterization of Amy1: An Alpha-Amylase from Cryptococcus flavus Expressed in Saccharomyces cerevisiae. Enzyme Res. 2011, 157294 (2011).

7. Gallo, R. L., Nizet, V., 2012. Endogenous production of antimicrobial peptides in innate immunity and human disease. Curr. Allergy Asthma Rep. 12, 427–435.

8. Giuliani, G. Pirri, and S. F. Nicoletto, “Antimicrobial peptides: an overview of a promising class of therapeutics,” Open Life Sciences, vol. 2, no. 1, pp. 31–33, 2007.

9. Gokhale, A. S., & Satyanarayanajois, S. (2014). Peptides and peptidomimetics as immunomodulators. Immunotherapy, 6(6), 755–774. https://doi.org/10.2217/imt.14.37

10. H. G. Boman, “Antibacterial peptides: key components needed in immunity,” Cell, vol. 65, no. 2, pp. 205–207, 1991.

11. H. Jenssen, C. D. Fjell, A. Cherkasov, and R. E. W. Hancock, “QSAR modeling and computer-aided design of antimicrobial peptides,” Journal of Peptide Science, vol. 14, no. 1, pp. 110–114, 2008.

12. H. Steiner, D. Hultmark, A. Engstrom, H. Bennich, and H. G. Boman, “Sequence and specificity of two antibacterial proteins involved in insect immunity,” Nature, vol. 292, no. 5820, pp. 246–248, 1981.

13. H.-S. Ahn, W. S. Cho, S.-H. Kang et al., “Design and synthesis of novel antimicrobial peptides on the basis of *α* helical domain of Tenecin 1, an insect defensin protein, and structure-activity relationship study,” Peptides, vol. 27, no. 4, pp. 640–648, 2006.

14. H.-T. Chou, T.-Y. Kuo, J.-C. Chiang et al., “Design and synthesis of cationic antimicrobial peptides with improved activity and selectivity against Vibrio spp,” International Journal of Antimicrobial Agents, vol. 32, no. 2, pp. 130–138, 2008.

15. Hancock, R. E. W., Sahl, H. G., 2016. Antimicrobial and host-defense peptides as new anti-infective therapeutic strategies. Nat. Biotechnol. 24, 1551–1557.

16. Hilchie, A. L., Wuerth, K. & Hancock, R. E. W. Immune modulation by multifaceted cationic host defense (antimicrobial) peptides. Nat. Chem. Biol. 9, 761–768 (2013).

17. J. M. Thomson and R. A. Bonomo, “The threat of antibiotic resistance in Gram-negative pathogenic bacteria: beta-lactams in peril!,” Current Opinion in Microbiology, vol. 8, no. 5, pp. 518–524, 2005.

18. K. V. R. Reddy, R. D. Yedery, and C. Aranha, “Antimicrobial peptides: premises and promises,” International Journal of Antimicrobial Agents, vol. 24, no. 6, pp. 536–547, 2004.

19. L. P. Miranda and P. F. Alewood, “Accelerated chemical synthesis of peptides and small proteins,” Proceedings of the National Academy of Sciences of the United States of America, vol. 96, no. 4, pp. 1181–1186, 1999.

20. Lindahl, E., Azuara, C., Koehl, P. & Delarue, M. NOMAD-Ref: visualization, deformation and refinement of macromolecular structures based on all-atom normal mode analysis. Nucleic Acids Res 34, W52–56, doi: 10.1093/nar/gkl082 (2006).

21. Liu, J., Cao, X. Cellular and molecular regulation of innate inflammatory responses. Cell Mol Immunol 13, 711–721 (2016). https://doi.org/10.1038/cmi.2016.58

22. M. A. Baker,W. L. Maloy, M. Zasloff, and L. S. Jacob, “Anticancer efficacy of Magainin2 and analogue peptides,” Cancer Research, vol. 53, no. 13, pp. 3052–3057, 1993.

23. M. Zasloff, “Antimicrobial peptides in health and disease,” The New England Journal of Medicine, vol. 347, no. 15, pp. 1199–1200, 2002.

24. Niyonsaba, F., et al., 2009. Antimicrobial peptides: key components of the innate immune system. Crit. Rev. Immunol. 29, 335–357.

25. Oddo A, Hansen PR. Hemolytic Activity of Antimicrobial Peptides. Methods Mol Biol. 2017;1548:427–435. doi: 10.1007/978-1-4939-6737-7_31. Erratum in: Methods Mol Biol. 2017;1548:E1. PMID: 28013523.

26. Pierce, B. G. et al. ZDOCK server: interactive docking prediction of protein-protein complexes and symmetric multimers. Bioinformatics 30, 1771–1773, doi: 10.1093/bioinformatics/btu097 (2014).

27. Pratiti Bhadra, Jielu Yan, Jinyan Li, Simon Fong, and Shirley W. I. Siu.* AmPEP: Sequence-based prediction of antimicrobial peptides using distribution patterns of amino acid properties and random forest. Scientific Reports, 1697 (2018).

28. R. E. W. Hancock and A. Patrzykat, “Clinical development of cationic antimicrobial peptides: from natural to novel antibiotics,” Current Drug Target—Infectious Disorders, vol. 2, no. 1, pp. 79–83, 2002.

29. R. Lai, Y.-T. Zheng, J.-H. Shen et al., “Antimicrobial peptides from skin secretions of Chinese red belly toad Bombina maxima,” Peptides, vol. 23, no. 3, pp. 427–435, 2002.

30. Roy, A., Kucukural, A. & Zhang, Y. I-TASSER: a unified platform for automated protein structure and function prediction. Nat Protoc 5, 725–738, doi: 10.1038/nprot.2010.5 (2010).

31. S. Haeberli, L. Kuhn-Nentwig, J. Schaller, and W. Nentwig, “Characterisation of antibacterial activity of peptides isolated from the venom of the spider Cupiennius salei (Araneae:Ctenidae),” Toxicon, vol. 38, no. 3, pp. 373–380, 2000.

32. Vidal-Limon A, Aguilar-Toalá JE, Liceaga AM. Integration of Molecular Docking Analysis and Molecular Dynamics Simulations for Studying Food Proteins and Bioactive Peptides. J Agric Food Chem. 2022 Feb 2;70(4):934–943. doi: 10.1021/acs.jafc.1c06110. Epub 2022 Jan 6. PMID: 34990125.

33. W. F. Broekaert, B. P. A. Cammue, M. F. C. De Bolle, K. Thevissen, G. W. De Samblanx, and R. W. Osborn, “Antimicrobial peptides from plants,” Critical Reviews in Plant Sciences, vol. 16, no. 3, pp. 297–323, 1997.

34. Wang, G., 2014. Human antimicrobial peptides and proteins. Pharmaceuticals 7, 545–594.

35. Wikler MA, Cockerill FR, Bush K, Dudley MN, Eliopoulos GM, Hardy DJ, et al. Methods for dilution antimicrobial susceptibility tests for bacteria that grow aerobically; approved standard—eighth edition Clinical and Laboratory Standards Institute; 2009

36. Y. Shai and Z. Oren, “From ‘carpet’ mechanism to de-novo designed diastereomeric cell-selective antimicrobial peptides,” Peptides, vol. 22, no. 10, pp. 1629–1641, 2001.

37. Yang, D., Biragyn, A., Kwak, L. W., Oppenheim, J. J., 2002. Mammalian defensins in immunity: more than just microbicidal. Trends Immunol. 23, 291–296.

38. Yang, L., Xie, X., Tu, Z. et al. The signal pathways and treatment of cytokine storm in COVID-19. Sig Transduct Target Ther 6, 255 (2021). https://doi.org/10.1038/s41392-021-00679-0

39. Z. Wang and G. Wang, “APD: the antimicrobial peptide database,” Nucleic Acids Research, vol. 32, supplement 1, pp. D590–D592, 2004.

40. Zasloff, M., 2002. Antimicrobial peptides of multicellular organisms. Nature 415, 389–395.

